# Motor control of the *Drosophila* antennae

**DOI:** 10.64898/2026.08.19.745841

**Authors:** Emily C. Kophs, Tobias C. McCabe, Sungwon Moon, Samuel Lee Sangmyung, Igor Siwanowicz, Marie P. Suver

## Abstract

Animals actively sense their surroundings to acquire behaviorally relevant environmental cues and stimuli. This dynamic acquisition of sensory information is enabled by active positioning of sensors and helps guide behavioral responses in dynamic environments. Yet how these active movements are controlled during behavior and coordinated with ongoing sensory acquisition is not fully understood. In the fruit fly *Drosophila melanogaster*, the antennae are crucial sensors for extracting important information from the environment including mechanosensory, olfactory, and auditory signals. Just four distinct muscles command movement of the antennae, providing a tractable model system for understanding efferent control of sensation. This work characterizes motor neurons used by *Drosophila* to actively position the antennae. We first identify antennal motor neurons in the central brain, and map each one from a comprehensive connectomic dataset to its peripheral muscle target. Our analysis of presynaptic inputs to the entire antennal motor system reveals a diverse array of premotor neurons for antennal motor control. We then provide genetic access to each motor unit by building a library of genetic lines with expression in antennal motor neurons. Using this library of antennal motor neuron lines, we next characterize motor unit function with quantitative behavior and optogenetics, revealing that the antennal motor system produces two primary movements in the dorsal-ventral and medial-lateral axes. Together, this work provides a comprehensive framework for understanding the motor control of an active sensor.

## Introduction

Animals actively move their sensory organs to acquire information from their environment. Examples of this can be found throughout the animal kingdom – whether it’s a rodent whisking to extract tactile information (Brecht et al. 1997; Diamond et al. 2008; Mitchinson et al. 2011; Staab et al. 2025), sniffing to refine olfactory stimulation (Crimaldi et al. 2022; Wachowiak 2011) or visual scanning (Toscani et al. 2013; Sakatani and Isa 2007; Cellini and Mongeau 2020; Fenk et al. 2010). Active sensor movements contribute to sensation for many behaviors that are essential for survival, such as during foraging, navigation, and predator avoidance, yet the neural mechanisms controlling these movements are not fully understood.

In insects, the antennae serve as multimodal active sensory organs, housing sensors for mechanosensation, olfaction, thermosensation, and hygrosensation. By actively positioning their antennae, insects can shape the sensory information they receive, allowing them to more effectively detect and respond to cues in their environment. For example, ants and locusts move their antennae to tune olfactory signals (Draft et al. 2018; Huston et al. 2015) and stick insects use antennal movements to probe for tactile information (Rajabi et al. 2018). But we still lack a comprehensive understanding of motor circuits controlling the antennae, limiting our ability to investigate how movement shapes sensory perception.

Using the fruit fly, *Drosophila melanogaster*, we can probe the neural circuits underlying the antennal sensory-motor system by leveraging the full brain connectome. We combine this with an expansive collection of genetic tools for labeling and manipulating small populations of neurons in behaving flies. Previous work identified the four muscles responsible for controlling antennal movements and described genetic driver lines targeting several of their motor neurons (Suver et al., 2023). However, the identity of these motor neurons in the brain, their specific muscle targets, and their functional roles in driving antennal movements remained unknown.

To establish a complete map of the antennal motor system, we identified each antennal motor neuron in the full brain connectome, matched individual neurons to their peripheral muscle targets, and identified genetic driver lines providing access to each motor unit. We analyzed the networks of presynaptic neurons for each motor neuron pair to understand the relative influence of major cell classes in the brain on antennal movement control. Next, we characterized the antennal movements elicited by their activation using optogenetics and quantitative behavioral analysis. Bridging across genetic, circuit, and behavioral analyses, this study reveals how the *Drosophila* antennal motor system is organized to control behavior.

## Results

### Four muscles innervated by five pairs of motor neurons control active antennal movements

To visualize the antennal motor system, we generated a three-dimensional reconstruction of the muscles within the antennae using confocal data (Figure 1, Video 1). Consistent with previous findings (Suver et al. 2023), the reconstruction delineates four muscle groups defined by their anterior-posterior position of insertion on the cuticle (numbered 1-4) within the scape. Muscles 1 and 2 appear to have one undivided muscle fascicle, whereas muscle 3 has three partitions connected to the cuticle via the same tendon (tripartite), and muscle 4 is bipartite.

**Figure 1.**
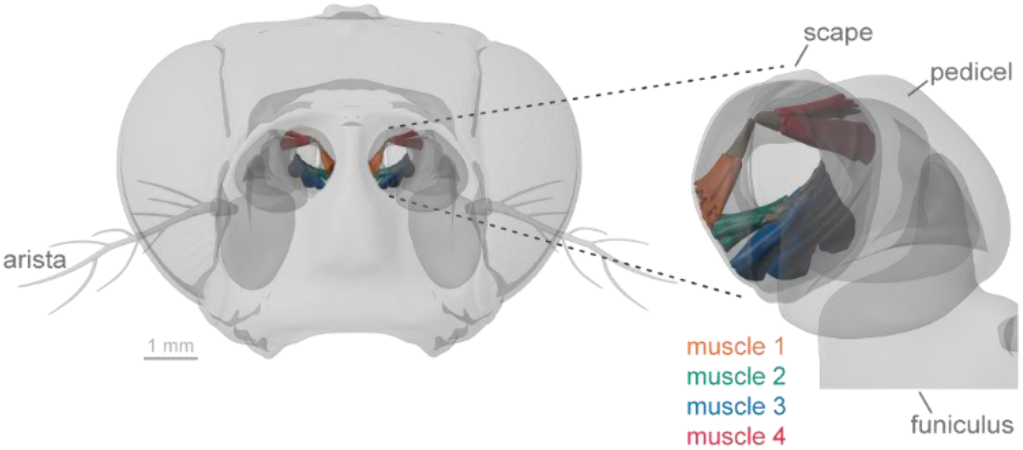
3D rendering of muscles in the *Drosophila* antennae. Left: Frontal view of the *Drosophila* head showing four muscles in the first segment of each antenna. Right: Single antenna enlarged to show the four muscles and their tendons in the first segment (scape), the second segment (pedicel) and part of the third segment (funiculus). Muscles are labeled 1-4, with 1 having the most anterior point of origin and 4 the most posterior.

To identify motor neurons innervating the four antennal muscles, we used publicly available whole brain connectomics data sets (FAFB: Dorkenwald et al. 2024; Schlegel et al. 2024; BANC: Bates et al. 2026; MCNS: Berg et al. 2025). All motor neurons innervating antennal muscles are predicted to be glutamatergic (Jan and Jan 1976), have a cell body and input processes in the brain, send an axonal projection through the antennal nerve, and ultimately synapse onto antennal muscles. Across all three connectomics data sets we examined, we identified seven pairs of neurons with cell bodies and input processes in the brain, and axons projecting out of the brain through the antennal nerve (Figure 2A, Table 1). We found that these seven pairs of neurons are the only efferent projections from the brain traveling through the antennal nerve in the three connectomic data sets. All putative motor neurons identified in this way are predicted to be glutamatergic (Eckstein et al. 2024), matching our expectation for motor neurons.

**Figure 2.**
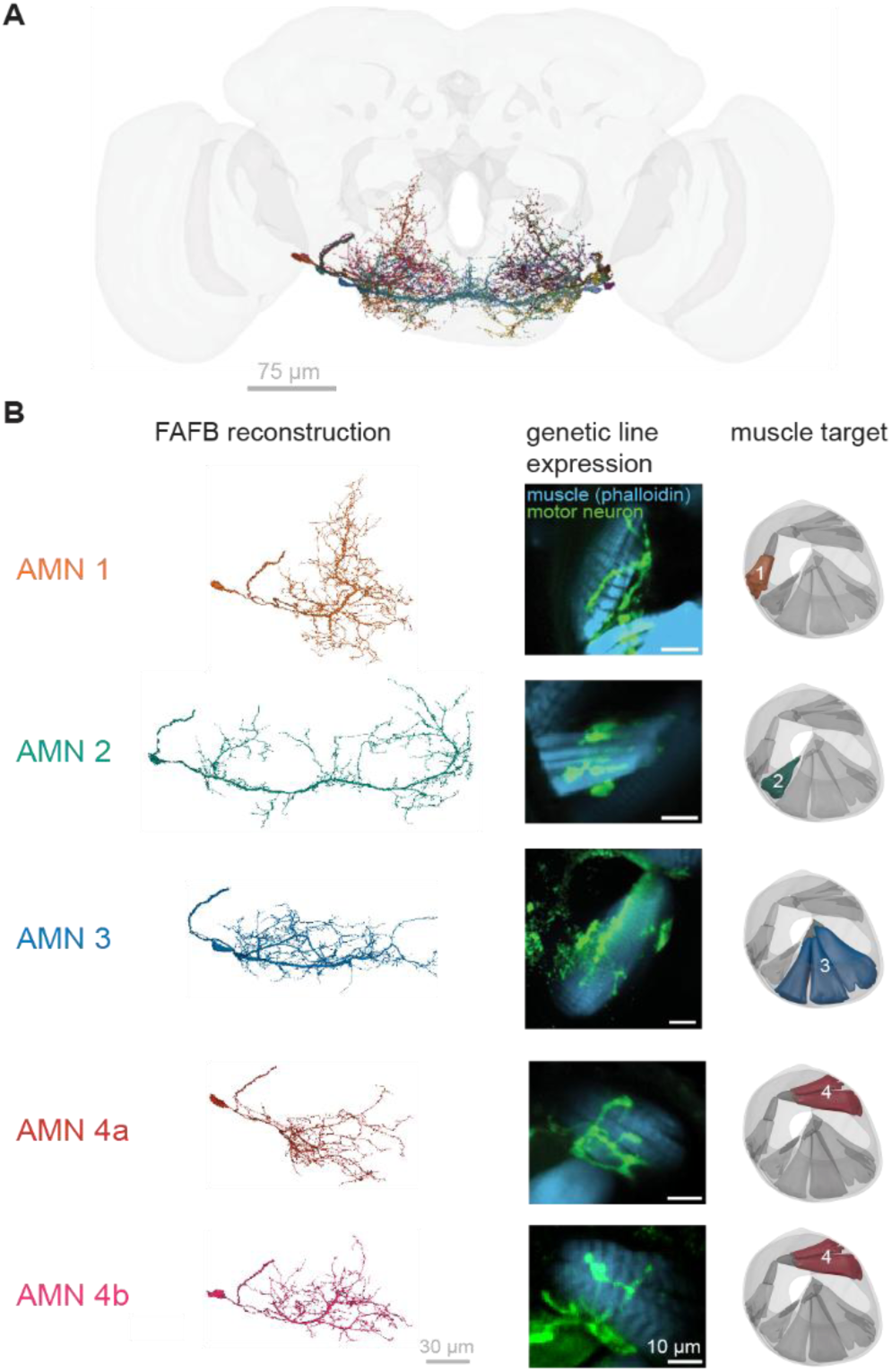
Five pairs of motor neurons innervate four muscles in the antennae. (A) Rendering of all five antennal motor neuron pairs in the female adult fly brain (FAFB; grey). (B) Mapping motor neurons in the connectome to their muscle targets in the antenna. Left: FAFB reconstructed motor neurons with respective names. Center: confocal images of each motor neuron expressing green fluorescent protein (green) synapsing onto its muscle target (blue). Right: innervated muscle schematic. Confocal images were obtained using *R18D07 > 20XUAS-SPARC2I Chrimson::tdTomato* for AMN 1, AMN 2, and AMN 4a, *R19H10 > UAS-Chrimson* for AMN 3, and *R75G05 > UAS-Chrimson* for AMN 4b.

**Table 1.**
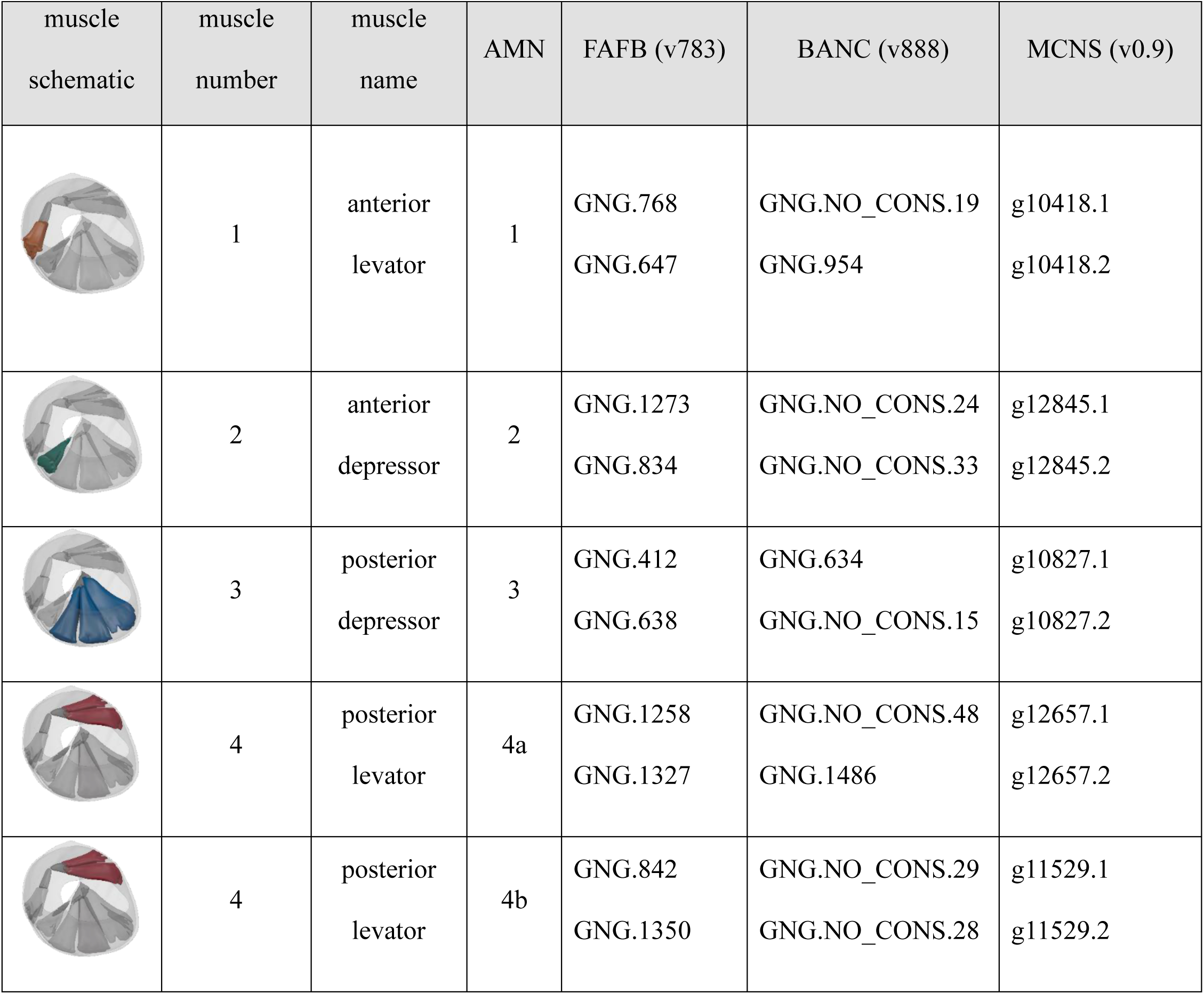
Mapping the antennal motor neurons from the connectome to their target muscles. The identities of the four antennal muscles with each corresponding antennal motor neuron are listed for the most current versions of the Female Adult Fly Brain (FAFB, Dorkenwald et al. 2024; Schlegel et al. 2024), Brain and Nerve Cord (BANC, Bates et al. 2026) and the Male Central Nervous System (MCNS, Berg et al. 2025).

| muscle<br>schematic | muscle<br>number | muscle<br>name | AMN | FAFB (v783) | BANC (v888) | MCNS (v0.9) |
| --- | --- | --- | --- | --- | --- | --- |
|  | 1 | anterior<br>levator | 1 | GNG.768<br>GNG.647 | GNG.NO_CONS.19<br>GNG.954 | g10418.1<br>g10418.2 |
|  | 2 | anterior<br>depressor | 2 | GNG.1273<br>GNG.834 | GNG.NO_CONS.24<br>GNG.NO_CONS.33 | g12845.1<br>g12845.2 |
|  | 3 | posterior<br>depressor | 3 | GNG.412<br>GNG.638 | GNG.634<br>GNG.NO_CONS.15 | g10827.1<br>g10827.2 |
|  | 4 | posterior<br>levator | 4a | GNG.1258<br>GNG.1327 | GNG.NO_CONS.48<br>GNG.1486 | g12657.1<br>g12657.2 |
|  | 4 | posterior<br>levator | 4b | GNG.842<br>GNG.1350 | GNG.NO_CONS.29<br>GNG.NO_CONS.28 | g11529.1<br>g11529.2 |

To verify that the motor neurons are clearly distinguishable from other cells innervating the antennae, we used the female adult fly brain (FAFB) whole brain EM reconstruction to investigate the size (area and volume) of all annotated cells in the antennal nerve. Our size analyses (Supplemental Fig. 1A, B) demonstrate that motor neurons, along with the bilateral pair of antennal campaniform sensilla, are the largest neurons in the antennal nerve, making them visually distinct from other cells. Further, because motor neurons are glutamatergic and campaniform sensilla are not, we leveraged genetic tools that label glutamatergic neurons. By imaging glutamatergic neurons labeled in the antennal nerve (*Vglut-GAL4>UAS-10xGFP*), we found that two axons leave the main branch of the antennal nerve through a proximal nerve that travels laterally (Supplemental Fig. 1C and 1D; Miller 1950). Five large axons continue through the antennal nerve towards the antenna (Supplemental Fig. 1E). In each of the three data sets, the two bilaterally symmetric neuron pairs that exit the brain via the antennal nerve had substantial presynaptic visual inputs in the inferior posterior slope (IPS); this prominent visual input to these two cells, but not the putative antennal motor neurons, suggests that these two cells may play an important role in visual sensation or behavior (Wei et al. 2020; Namiki et al. 2018). By investigating the projection patterns of a genetic line labeling one eye motor neuron and one putative antennal motor neuron identified in a previous study (Fenk et al. 2022), we found that one axon travels straight through the antennal nerve towards the antennae and ultimately synapses onto an antennal muscle, whereas the other axon travels through the proximal nerve towards a retinal eye muscle. This result further supported our conclusion that the two large glutamatergic neurons exiting the antennal nerve are the two retinal eye motor neurons (Supplemental Fig. 1C). Thus, we concluded that the five remaining pairs of large efferent neurons in the antennal nerve were very likely antennal motor neurons (AMNs, Figure 2). Although the individual antennal motor neurons within each identified left/right pair are bilaterally symmetrical, the different pairs have distinct projection patterns within the gnathal ganglia (Figure 2B).

We used anatomical tools linked to connectomic resources to identify potential driver lines targeting the five antennal motor neuron pairs (see Methods; Clements et al. 2024; Azevedo et al. 2024). We then systematically examined the expression pattern of each candidate line in the antennae to determine whether a labeled neuron directly innervated the antennal muscles.

Through this screening process, we generated a library of driver lines targeting individual and subsets of AMN pairs (Table 2). After identifying driver lines that label neurons innervating specific antennal muscles, we then sought to match each muscle to its corresponding motor neuron(s) in the central brain. Stochastically limiting fluorescent effector expression in each fly (using SPARC, <u>Isaacman-Beck et al. 2020)</u> enabled us to clearly image individual motor neurons in the brain and their muscle targets in the antennae. Using these images, we matched each AMN pair in the connectome to an antennal muscle pair and the driver lines to access them (Figure 2B). We found that each muscle is innervated by one motor neuron except for muscle four; additional confocal imaging data revealed two distinct axons innervating muscle 4, indicating that this muscle is controlled by two separate AMNs (Supplemental Figure 2A). Each of the two motor neurons that synapse onto muscle 4 appear to innervate both muscle partitions (Supplemental Figure 2B, C). Similarly, the motor neuron synapsing onto muscle 3 innervates all three muscle partitions (data not shown). We named each of the AMN pairs according to the muscle they target: AMN 1 innervates muscle 1, AMN 2 innervates muscle 2, AMN 3 innervates muscle 3, and AMN 4a and 4b both innervate muscle 4.

**Table 2.** A library of driver lines with expression in antennal motor neurons. Our collection of GAL4 and split-GAL4 lines provides genetic access to individual AMNs, groups of AMNs, and one antennal muscle. We also note off-target expression in the antennal nerve. We classified any line in which we observed neurons in the antenna that were not innervating muscles as having off-target sensory expression.

| Driver Line | Muscle Number(s) | Muscle Name(s) | Motor Neuron(s) | Off-Target Expression |  |
| --- | --- | --- | --- | --- | --- |
|  |  |  |  | Eye motor neurons | Sensory neurons |
| *VT028320-GAL4* | 1, 3 | anterior levator, posterior depressor | AMN 1, AMN 3 | ☒ | ☐ |
| *VT045575AD;VT026387DBD* | 2 | anterior depressor | AMN 2 | ☐ | ☐ |
| *R74C10-GAL4* | 3 | posterior depressor | N/A | ☐ | ☐ |
| *SS40851* | 3 | posterior depressor | N/A | ☐ | ☒ |
| *R55E06-GAL4* | 4 | posterior levator | AMN 4a | ☐ | ☒ |
| *R75G05-GAL4* | 4 | posterior levator | AMN 4b | ☐ | ☐ |
| *VT012411AD;R10B11DBD* | 4 | posterior levator | AMN 4b | ☒ | ☐ |
| *R19H10-GAL4* | 2(stochastic), 3 | anterior depressor, posterior depressor | AMN 2 (stochastic)<br>AMN 3 | ☒ | ☒ |
| *R18D07-GAL4* | 1(stochastic), 2, 4 | anterior levator, anterior depressor, posterior levator | AMN 1 (stochastic)<br>AMN 2<br>AMN 4a<br>AMN 4b | ☒ | ☐ |
| *R91F02-GAL4* | 2(stochastic), 3, 4 | anterior depressor, posterior depressor, posterior levator | AMN 2 (stochastic)<br>AMN 3<br>AMN 4a or b | ☐ | ☒ |

### Shared upstream connectivity reveals functional groupings

To better understand how each motor neuron is recruited for behavior, we examined the number and identity of presynaptic partners for each antennal motor neuron (Figure 3). Using data from the adult fly brain (FAFB) connectome (Dorkenwald et al. 2024; Schlegel et al. 2024), we first generated a network map of the neurons upstream of each of the AMNs (Figure 3A). As a population, the AMNs received just over 800 unique presynaptic inputs. Individual AMNs received between 67 and 208 total presynaptic partners, with the bilateral pair AMN 3 together receiving the largest number of inputs (399 total). The distance between each AMN in the directed presynaptic network (Figure 3A) reflects the number of shared presynaptic partners for each AMN (i.e., AMN nodes that are closer together have more shared input partners, whereas farther-spaced nodes indicate more distinct presynaptic inputs).

**Figure 3.**
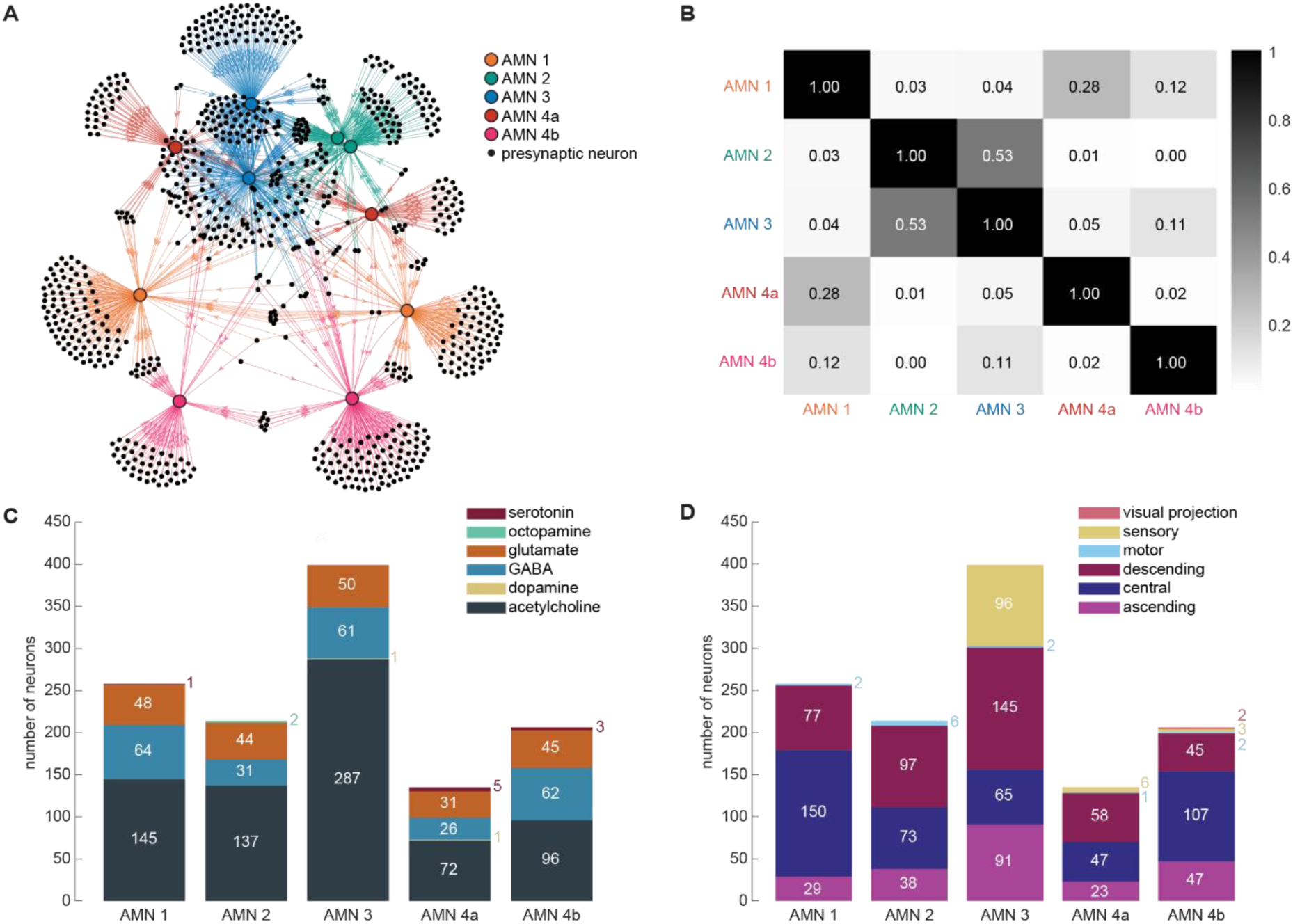
Connectomic analysis provides insight into the function of motor neuron pairs. (A) Connectivity network of AMNs and their presynaptic partners. Colorful circles represent AMNs, and small black dots represent presynaptic neurons. Arrows indicate direction of the connection between presynaptic partners and the AMNs. (B) Weighted cosine similarity matrix of AMN presynaptic networks. Each AMN pair is represented on the x and y axes. Higher values indicate a greater similarity in presynaptic partners. (C) Neurotransmitter distribution across AMN inputs. Each bar represents the total number of presynaptic neurons for an AMN pair. Colored segments indicate the number of presynaptic partners belonging to each predicted neurotransmitter class. (D) Superclass distribution across AMN input partners. Each bar represents the total number of presynaptic neurons for an AMN pair. Colored segments represent the number of presynaptic partners belonging to different predicted super classes.

To quantify shared versus distinct presynaptic parters amongst AMN pairs, we calculated pairwise cosine similarity scores based on their presynaptic inputs and created a cosine similarity matrix (Figure 3B). This matrix revealed two groups of motor neurons with similar presynaptic influences. AMN 1 has highest similarity scores with AMN 4a (0.28), and AMN 4b (0.12), which are relatively high when compared to AMN2 (0.03) and AMN3 (0.04). Likewise, AMN 2 and AMN 3 share the highest cosine similarity score with one another (0.53). We also observed very low similarity scores between the two motor neurons innervating muscle 4 (0.02).

We next classified presynaptic input into each AMN pair by neurotransmitter type. The stacked bar plots In Figure 3C show the total number of presynaptic inputs to each AMN pair broken down by predicted neurotransmitter type. We found that direct input from modulatory neurons (classified as serotonin, octopamine, or dopamine neurons) to any of the motor neuron pairs was sparse, making up just 1.47% of the total input across all the motor neurons. Most inputs were either excitatory or inhibitory, with each antennal motor neuron pair receiving a mix of both types. Excitatory (cholinergic) inputs were the largest group represented in the presynaptic network.

We also characterized the presynaptic inputs to each AMN pair according to their superclass designation in the FAFB dataset (Figure 3D). The sensory neurons projecting directly to the AMNs are entirely mechanosensory – no visual, olfactory, or thermosensory neurons appear to directly project to the AMNs. The mechanosensory inputs include Johnston’s Organ Neuron (JONs), bristles, and the singular bilateral pair of antennal campaniform sensilla. (One of the inputs does not have a defined cell type in FAFB, but based on morphology, is likely a JON.) Only three AMNs receive sensory input, and most of these synapses (96/105 total) are onto AMN 3. As a group, the antennal motor neurons receive roughly equal input from JONs and bristles, both as measured by number of neurons (59 JONs versus 44 bristle neurons) and synapses (607 JON synapses versus 783 bristle synapses) in the FAFB data set. Just over half of the bristle neurons providing input originate on the antennae (27/44, roughly 60%); other bristle inputs are likely from frontal, ocellar, orbital, or postocellar bristles (Eichler et al. 2024). The single antennal campaniform sensilla synapsing onto AMN 3 provides relatively more modest synaptic input, with 28 detected synapses across this bilateral pair of motor neurons. A small number of motor neurons appear to exist in the antennal premotor network as well (between 0 and 4 per antennal motor neuron, 13 total across the entire network), which can be attributed to connectivity between antennal motor neurons, in addition to inputs from the two eye motor neurons, a neck motor neuron, and one proboscis motor neuron.

### Optogenetic activation of motor neurons drives elevation and depression of antennae

To determine the functional role of each motor neuron and muscle group, we optogenetically activated tethered flies from each of the driver lines in our library (Figure 4A). We recorded the antennal response to this activation with two orthogonal cameras to capture changes in antennal position along both the lateral and dorsal axes (Figure 4B-C).

**Figure 4.**
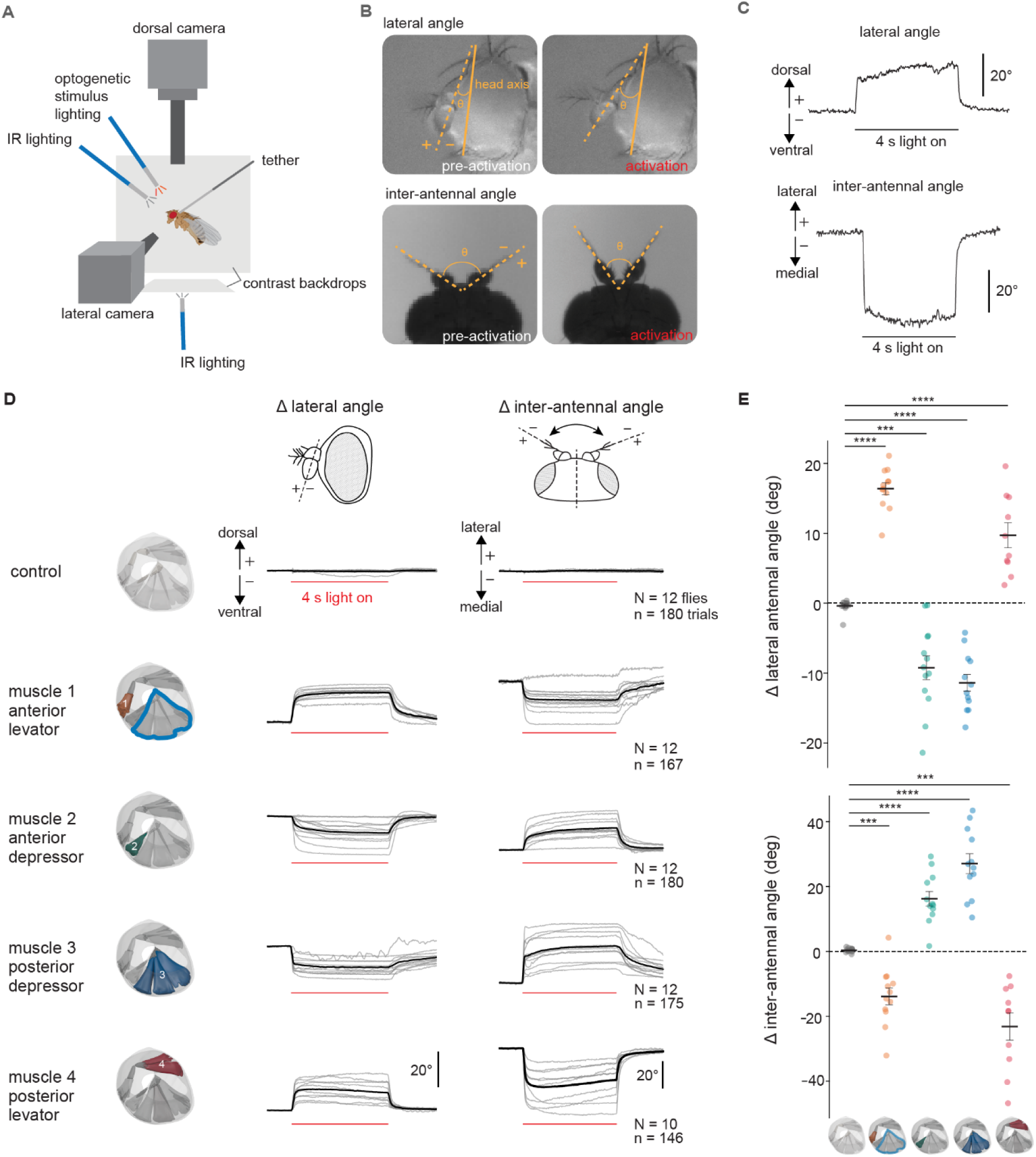
Two functional pairs of antennal muscles – the levators and depressors – guide antennal movements. (A) Schematic of the optogenetic behavioral apparatus. The fly was illuminated with infrared light and antennal movements were captured by dorsal and lateral cameras. (B) Top: example video frames from the lateral camera view before (left) and during the activation stimulus (right). Ventral antennal movements are negative, and dorsal movements are positive. Bottom: example video frames from the dorsal camera view before the activation stimulus (left) and during the activation stimulus (right). Antennal movements towards the body midline are negative and movements away from the body midline are positive. (C) Top: Example single trial trace representing dorsal-ventral antennal movements during motor neuron activation. Bottom: Example single-trial trace of lateral-medial antennal movements during motor neuron activation. (D) Left: Schematics representing targeted muscles. Outlined muscle indicates known off-target expression in addition to muscle of interest. Right: Comparison of antennal movements in response to the activation stimulus in control flies (*Canton-S > UAS-Chrimson*), flies with AMN 1 and AMN 3 labeled (*VT028320 > UAS-Chrimson*), flies with AMN 2 labeled (*VT045575AD;VT026387DBD > UAS-Chrimson*), flies with muscle 3 labeled (*R74C10 > UAS-Chrimson*), and flies with AMN 4a labeled (*R55E06 > UAS-Chrimson*). Grey lines represent the average antennal response of individual flies and black traces represent the average response across flies. (E) The change in antennal was quantified by subtracting the average pre-stimulus position from the average position during the last second of the stimulus. Each colorful dot represents one fly. Black bars represent the mean ± SEM. In comparison to the control, activation of AMN 1 and AMN 3 elevated the antennae (Mann-Whitney U; p = 3.658 × 10⁻⁵) and brought it towards the midline (t-test, p = 0.0002). Activation of AMN 2 pulled the antennae down (Mann-Whitney U, p = 0.0002) and away from the midline (t-test, p = 1.820 × 10⁻⁵)). Activation of muscle 3 brought the antennae down (Mann-Whitney U, p = 3.658× 10⁻⁵) and away from the midline (t-test, p = 2.727× 10⁻^6^). Activation of AMN 4a brought the antennae up (Mann-Whitney U, p = 8.734× 10⁻^5^) and toward the midline (t-test, p = 0.0004).

We were unable to find driver lines that selectively labeled AMN 1 and AMN 3 without additional expression in other AMNs. To address this, we used one driver line labeling muscle 3 (with no AMNs labeled, Suver et al. 2023), and compared activation responses in this genotype compared to another line labeling both AMN 1 and AMN 3. This enabled us to account for the contribution of muscle 3 (via AMN 3) to antennal movements when analyzing the combined activation, thereby indirectly determining the movements elicited by AMN 1.

Activation of the driver line labeling AMN 1 together with AMN 3 (*VT028320-GAL4*, Figure 4D) drove the antennae dorsally (Mann-Whitney U; p = 3.658 × 10⁻⁵) and inwards toward the midline (medially, t-test, p = 0.0002). Activation of AMN 2 (*VT045575-AD*; *VT026387-DBD*, Figure 4D) pulled the antennae ventrally (Mann-Whitney U, p = 0.0002) and laterally, away from the midline (t-test, p = 1.820 × 10⁻⁵). Activation of muscle 3 alone also significantly depressed the antennae (Mann-Whitney U, p = 3.658× 10⁻⁵) and pulled them away from the midline (t-test, p = 2.727× 10^⁻6^). Finally, activation of AMN 4a (*R55E06-GAL4*; Figure 4D) significantly elevated the antennae dorsally (Mann-Whitney U, p = 8.734× 10⁻5) while bringing them toward the midline (t-test, p = 0.0004). In contrast, activation of AMN 4b using two independent driver lines did not elicit a consistent antennal response (*R75G05-GAL4* and *VT012411-AD*; *R10B11-DBD;* Supplemental Fig. 3). Although some flies exhibited the same upward and inward movement observed with AMN 4a activation, other individuals displayed no detectable change in antennal position.

Following this quantification, we named each muscle according to both its anterior-posterior position within the first antennal segment and the movement elicited during activation: anterior levator (muscle 1), anterior depressor (muscle 2), posterior depressor (muscle 3), and the posterior levator (muscle 4).

## Discussion

In this study, we set out to comprehensively characterize the motor system of the *Drosophila* antenna. With only four muscles and five motor neurons, the *Drosophila* antenna offers a remarkably tractable motor system that is small enough to permit genetic, connectomic, and functional characterization of every component. This simplicity provides a unique opportunity to investigate how neural circuits coordinate movement of an active sensory organ. We generated a 3D reconstruction of the antenna, including each muscle and its associated tendon (Figure 1), identified every antennal motor neuron in a connectomic data set and matched it to its muscle target, and established a library of genetic driver lines that provides access to individual motor neurons or defined motor neuron subsets (Figure 2, Table 2). Next, our connectomic analyses revealed distinct patterns of presynaptic partners onto each motor neuron pair, enabling us to generate testable hypotheses about how different antennal movements are recruited during behavior (Figure 3). Finally, using targeted activation of muscles and motor neurons, we quantified the antennal movements produced by each motor unit (Figure 4).

Previous studies established *Drosophila* as a powerful model for linking neural circuits to behavior through characterization of multiple motor systems, including those controlling the legs (Azevedo et al. 2020), proboscis (McKellar et al. 2020), neck (Gorko et al. 2024), halteres (Dickerson et al. 2019), and flight muscles (Lindsay et al. 2017). Here, we add the antennal motor system to this growing collection of characterized motor circuits. We also link three previously separate resources: the central nervous system connectome, the peripheral musculature of an active sensor, and genetic access to the motor neurons that drive antennal movement (Figure 2, Table 2).

Our work revealed that of the four antennal muscles, muscle 4 is uniquely innervated by two motor neurons (Supplement Fig. 2), whereas each of the other three antennal muscles are innervated by a single motor neuron (Figure 2B). Optogenetic activation of either motor neuron innervating muscle 4 individually produces antennal elevation (Figure 4), this supports the idea that both neurons can drive contraction of this muscle. Yet the presynaptic inputs to AMN 4a and AMN 4b were dramatically non-overlapping, hinting at different functions. In the *Drosophila* leg motor system, muscles receiving input from multiple motor neurons often contain distinct fast- and slow-twitch motor units that expand the dynamic range of force production (Azevedo et al. 2020). This contrasts with the wing and haltere motor systems, where just one motor neuron controls each muscle (except for the tp1 muscle, which stochastically receives one or two motor neuron inputs; Cheong et al. 2026; Teoh et al. 2025). Modulatory neurons also innervate *Drosophila* skeletal muscle (Pauls et al. 2018), raising the possibility that one of the motor neurons innervating antennal muscle 4 serves a specialized role in modulating activity of the motor unit. Whether these two motor neurons serve analogous physiological roles or instead support distinct modes of antennal motor control, remains an important question for future functional studies.

A map of presynaptic network connections to antennal motor neurons (Figure 3A) revealed that the two contralaterally projecting motor neuron pairs, AMN 2 and AMN 3, share the greatest proportion of presynaptic partners with their corresponding mirror neurons. This observation complements the distinctive morphology of these cells, with each neuron’s projections crossing contralaterally across the midline of the brain. In addition, our cosine similarity analysis points towards AMN2 and AMN3 as having the greatest number of shared presynaptic partners out of the entire set of antennal motor neurons. Together, these findings suggest that the bilateral AMN2 and AMN3 pairs are recruited by common upstream circuits, allowing simultaneous activation of homologous depressor muscles in both antennae.

Our cosine similarity analysis (Figure 3B) further suggests that the antennal motor neuron pairs may be organized into functional groups according to the movements they produce. The three motor neuron pairs innervating levator muscles (muscles 1 and 4) exhibited greater cosine similarity to one another than to the two pairs innervating depressor muscles. Likewise, the depressor motor neuron pairs (muscles 2 and 3) were more similar to each other than to the levator motor neurons. This organization suggests that upstream circuits are structured around movement classes rather than individual muscles, with common networks recruiting muscles that produce similar actions.

One particularly striking observation from our connectomic analysis the antennal motor neuron presynaptic network was a substantial amount of sensory input, but only from mechanosensory neurons (Figure 3D). Previous work in *Drosophila* demonstrated a role for visual influence on antennal movements (Mamiya et al., 2011; Mills et al., 2026), so our work suggests that this visual influence is indirect. Substantial synaptic input to the antennal motor neurons also comes via descending neurons (Figure 3D), offering one putative source for indirect visual input. The direct sensory input to antennal motor neurons originates from exclusively mechanosensory neurons, including Johnston’s organ neurons, bristles, and the antennal campaniform sensilla (Figure 3D), indicating that fast antennal movements may be primarily dictated by these mechanosensory pathways. Rapid sensorimotor feedback could enable antennal position to be adjusted in response to mechanical perturbations, as previously shown in hawkmoths (Krishnan et al., 2012, Natesan et al., 2019). Surprisingly, however, feedback appears to not arise exclusively from mechanosensory bristles, which have been shown to synapse onto or near antennal motor neuron processes in other insects (Goldammer & Durr 2018; Krishnan et al. 2012). Instead, we observed roughly equal input from JONs and bristles, and additional direct input from the antennal campaniform sensilla. Whether this diversity of direct mechanosensory inputs represent a conserved feature across insects or a specialization in fruit flies remains an interesting question to address in future behavioral and physiological studies.

Overall, our optogenetic activation experiments revealed that one pair of muscles elevates the antenna and the other pair depresses it. This suggests that the antennal motor system is organized into functionally opposing muscle groups, consistent with previous work in stick insects (Dürr et al. 2001; Dirks and Dürr 2011), hawkmoths (Sant and Sane 2018), and cockroaches (Okada and Toh 2004). Although we observed some variation in medial-lateral movements between muscles within each functional group, these differences were relatively small compared to their similar movements in the dorsal-ventral axis, and they matched in sign for each group. Thus, we are confident in grouping the motor neurons into two distinct classes, one that moves the antennae dorsally (elevates with respect to the longitudinal axis of the animal) and medially (towards the midline), and another that positions the antennae ventrally (depressors) and somewhat more laterally (away from the midline).

Our muscle characterization allows us to generate hypotheses about how individual muscles contribute to specific behaviors. During flight, flies and other insects characteristically elevate their antennae upwards (dorsally) and draw it in towards the midline (Mills et al. 2026; Gewecke 1970; Mamiya et al. 2011; Staudacher et al. 2005; Krishnan et al. 2012; Heran 1959). In this study, we found that muscles 1 and 4 both elevate the antennae and draw the antennae inwards towards the midline, suggesting that one or both may contribute to flight-induced antennal positioning. Activation of a genetic driver line that targets both muscle 2 and muscle 4 (*R18D07-GAL4)* reproduced the elevated and medialized antennal flight positioning in non-flying flies (Supplemental Figure 3). This result suggests that the two muscles contribute to different components of antennal posture: levator activity elevates the antenna along the dorsal-ventral axis, whereas depressor activity contributes to medial-lateral positioning. This interpretation is consistent with observations in the stick insect, where contraction of antennal depressor muscles has been shown to contribute to both medial and lateral antennal movements (Dürr et al. 2001), demonstrating that depressor muscles can play a broader role in shaping antennal posture than their name alone would imply. This complete motor map provides a foundation for understanding how antennal movements are controlled across behaviors and, ultimately, how active control of a sensory organ shapes sensory acquisition by the nervous system.

## Methods

**Table 2.** Reagents and fly lines used in this study.

| Reagent or Resource | Source | Dilution | Identifier |
| --- | --- | --- | --- |
| **Antibodies** |  |  |  |
| Mouse anti-bruchpilot | Developmental Studies Hybridoma Bank | 1:8 | nc82-s |
| Alexa Fluor 488 Phalloidin | Invitrogen | 1:40 | A12380 |
| Rabbit anti-DsRed | Takara Bio | 1:500 | 632496 |
| Rabbit anti-GFP | Invitrogen | 1:50 | A11122 |
| Alexa633-conjugated goat anti-mouse | Invitrogen | 1:250 | A21052 |
| Alexa568-conjugated goat anti-rabbit | Invitrogen | 1:250 | A11011 |
| Alexa488-conjugated goat anti-rabbit | Invitrogen | 1:250 | A11039 |
| **Experimental models: Organisms/strains** |  |  |  |
| Canton-S | Dickinson Lab |  | N/A |
| *+(HCS); +; P{10xUAS-IVS-SYR1-GFP-p10} attP2* | Pfeiffer et al 2012 via the Dickinson Lab, Caltech |  |  |
| *w[1118]; P{y[+t7.7] w[+mC]=20XUAS-IVS-CsChrimson.mVenus}attP40* | BDSC |  | RRID: BDSC_55135 |
| *20XUAS-SPARC2D Chrimson::tdTomato* | BDSC |  | RRID: BDSC_84143 |
| *20XUAS-SPARC2I Chrimson::tdTomato* | BDSC |  | RRID: BDSC_84144 |
| *PhiC31; T(2,3)engrailed/CyO; TM3,ser* | Nagel Lab, NYU Langone |  | N/A |
| *R18D07 GAL4* | BDSC |  | RRID: BDSC_48813 |
| *R91F02 GAL4* | BDSC |  | RRID: BDSC_40584 |
| *R74C10 GAL4* | BDSC |  | RRID: BDSC_39848 |
| *SS40851* | BDSC |  | RRID: BDSC_88411 |
| *R55E06 GAL4* | BDSC |  | RRID: BDSC_39121 |
| *R75G05 GAL4* | BDSC |  | RRID: BDSC_47958 |
| *R19H10 GAL4* | BDSC |  | RRID: BDSC_48868 |
| *VT028320 GAL4* | KDRC |  | KDRP013467 |
| *VT012411-AD;R10B11-DBD* | This study |  | RRID: BDSC_72280<br>and RRID:<br>BDSC_69860 |
| *VT045575-AD;VT026387-DBD* | This study |  | RRID: BDSC_72072<br>and RRID:<br>BDSC_73993 |
| *PhiC31; +/-cyo; R18D07*<br>*GAL4/ser* | This study |  |  |
| *PhiC31; +/-cyo; R55E06*<br>*GAL4/ser* | This study |  |  |
| *PhiC31; +/-cyo; R75G05*<br>*GAL4/ser* | This study |  |  |
| *PhiC31; +/-cyo; R19H10*<br>*GAL4/ser* | This study |  |  |
| *PhiC31; VT012411AD/cyo;*<br>*R10B11DBD GAL4/ser* | This study |  |  |
| *vGlut-GAL4* | BDSC |  | RRID: BDSC_24635 |

### Fly stocks and rearing

We used adult female *Drosophila melanogaster* between 3 and 10 days old for all experiments. Flies were raised on standard cornmeal-molasses food at 25°C in a 12-hour light/dark cycle. For optogenetics experiments, adult flies were transferred to food enriched with all-trans retinal and kept in light-shielded vials for 24-72 hours prior to the experiments. To prepare the all-trans retinal food, 50 μl of 35 mM all-trans retinal (Sigma-Aldrich R2500-100MG dissolved in ethanol) was added to 0.75 grams of rehydrated potato flakes and placed on top of the standard cornmeal-molasses food.

### Generating data set for creating the fly model

Flies were anesthetized on ice, briefly washed with ethanol and dissected under PBS-T (PBS + 0.1% Triton X-100). Heads, wings, thoraces with abdomens, fore-, mid- and hind legs belonging to the same fly to were transferred to individual tubes. All body parts except the wings were incubated with 0.25 mg/ml trypsin in PBS-T for 48 hours at 37°C to remove the soft tissues. The cuticle was then bleached in 20% H2O2 for 24h, dehydrated in ethanol and mounted in methyl salicylate (Sigma-Aldrich #M6752).

Serial optical sections were obtained at 1 µm with a LD-LCI 25x/0.8 NA objective, and at 0.3 µm with a Plan-Apochromat 40x/0.8 NA objective. Green autofluorescence characteristic for hard/sclerotized cuticle was excited by a 488 nm laser.

### Constructing the model

3D meshes were extracted from the confocal stacks using Fiji’s 3D viewer plugin (http://fiji.sc/) and imported into Blender (http://www.blender.org.). A 3D model was constructed from meshes representing the head, thorax/abdomen, wing and fore-, mid- and hind-leg of a single female fly, as described in Vaxenburg et al. 2025. Appendage meshes were mirrored across the body’s medial plane.

### Screening and Selection of Driver Lines

To find driver lines targeting antennal motor neurons, we started by identifying the antennal motor neurons in the Flywire Brain Dataset (FAFB v783). The five pairs of these cells are classified as motor neurons with a process leaving the brain through the antennal nerve. We matched each of these cells to candidate driver lines using the ColorMIP matching widget on BrainCircuits.io and the matching tool on NeuronBridge (Clements et al. 2024; Azevedo et al. 2024). We narrowed this list of potential driver lines by carefully scrutinizing the confocal brain images available for each driver line on FlyLight (Dionne et al. 2018; Jenett et al. 2012; Tirian and Dickson 2017). We then obtained lines with evidence of convincing cell morphology matching antennal motor neurons from the Bloomington Drosophila Stock Center. We crossed these lines to a *10XUAS-GFP* reporter for visualization of fluorescence in the antennal nerve. To do this, we used UV glue (Wayin UV Resin Hard) to immobilize live flies in a custom press-fit holder and then dissected off a small piece of cuticle on the top of the head capsule, directly above the first antennal segments, to expose the antennal nerves. We then examined the antennal nerves for fluorescence using a fluorescent LED (Sutter Instruments LAMBDA HPX-L5). This step allowed us to shortlist only driver lines with visible fluorescence in the antennal nerves for further dissections. For these lines, we proceeded with our brain and antennae dissection and imaging protocols. Only driver lines with clear neuron innervation onto muscles in the antennae were added to our library.

### Immunohistochemistry and Imaging

We submerged flies in 70% ethanol for 30 seconds to remove lipids from the cuticle, then transferred them to a dissecting dish with 1X PBS (phosphate-buffered saline, Corning MT21040CV). We then dissected the brain out of the head capsule, taking care not to sever the antennal nerves, keeping the antennae attached to the brain. We transferred each brain-antennae complex to 4% paraformaldehyde (Electron Microscopy Sciences 15710, diluted in PBS) and fixed them for 20 minutes in a light-proof box at room temperature. After washing 3X in PBS with 0.2% triton-x (Santa Cruz Biotechnology sc-29112) we incubated the brain-antennae complexes in 5% normal goat serum in PBST (Vector Laboratories, S-1000-20) for 20 minutes at room temperature. For staining, we incubated each brain-antennae complex in primary antibody solution (see Table 2 above) overnight at room temperature, washed 3X in PBST, and then incubated again overnight in secondary antibody solution before a final 3X wash in PBST (see Table 2 above). To mount both the antennae and the brain flat on a coverslip, we carefully severed the antennal nerves to separate the brain from the antennae just before mounting. We then flipped the coverslips onto a slide with vectashield (Vector Laboratories H100010) and sealed it with nail polish. We performed all imaging on a Zeiss LSM 880 confocal with a Plan-Apochromat 20x/0.80NA objective.

### Optogenetic Experiments

To capture movements of the antennae, we used a camera mounted dorsally relative to the fly (Allied Vision Guppy Pro F-031 with InfiniStix 68mm/1.00x lens) and another camera mounted laterally to capture a view of the left antennae (Allied Vision Guppy Pro F-031 with InfiniStix 44mm/3.00x lens). The fly was illuminated from below with infrared light (850nm, ORDRO LN-3 Studio IR Light Accessory). A fiber optic (ThorLabs M118L02) was positioned directly in front of the fly to deliver the activation stimulus. For activation, we used a 590 nm LED (ThorLabs M590F3) at an intensity strong enough to elicit an antennal response across all driver lines (0.697 mW or 9.87 mW/cm^2). Each fly experienced 20 randomized trial conditions, 15 with an activation stimulus and 5 without. Trials were 10 seconds long, with 2 seconds of no stimulus (light off) followed by 4 seconds with the stimulus light on and a final 4 seconds of no stimulus. We cold-anesthetized the flies and dissected their legs at the femoral-trochanter joint to prevent grooming of the antennae. We used ultraviolet curing glue (Topchase UV Resin) to fix the head to the thorax and to attach a tungsten (0.005’’ diameter, AM Systems 716000) pin tether to the thorax.

To track antennal movements, we used DeepLabCut (version 2.3.6). We trained our dorsal network on 340 randomly selected frames from 17 different flies across genotypes. For this network, we tracked 8 points on the head of the fly, including 2 static points used to determine an axis relative to the fixed head. Our lateral network was trained on 525 randomly selected frames from 35 different flies across genotypes. For this network, we tracked 6 points on the antenna and 2 static points on the head capsule. We used a ResNet-50 neural network, trained over 500,000 iterations with a training-to-test fraction of 0.8. The dorsal network yielded a test error of 1.71 pixels and a train error of 1.66 pixels in an image size of 640×347 pixels. The lateral network yielded a test error of 2.68 pixels and a train error of 2.24 pixels in an image size of 640×347 pixels.

### Statistical Analysis

All data was collected using MATLAB 2024b and analyzed using custom python scripts. Alpha values were set to 0.05 for statistical tests, with * representing ≤0.05, ** representing ≤0.01, *** representing ≤0.001, and **** representing ≤0.0001. We used Shapiro-Wilk’s test to test the normality of the data (python: scipy.stats.shapiro) and Levene’s test for equal variance (scipy.stats.levene). Lateral antennal angles were computed by measuring the angle between the funiculus and a stable head axis for each fly. Inter-antennal angles were computed by adding the angle between the left antenna and the midline of the head to the angle between the right antenna and the midline of the head. We compared the average angle changes in each experimental group to those of the control group. For normally distributed data, we used independent t-tests (python: scipy.stats.ttest_ind). For data that was not normally distributed, we used the Mann-Whitney U test (python: scipy.stats.mannwhitneyu).

### Connectomic Analyses

We used custom MATLAB (2024b) scripts to analyze data from version 783 of the Female Adult Fly Brain connectome (FAFB, Dorkenwald et al. 2024; Schlegel et al. 2024). We downloaded the FAFB data (v783) from Codex: https://codex.flywire.ai/api/download?dataset=fafb.

Using their unique cell IDs, we extracted the presynaptic input data for each of the ten antennal motor neurons into separate CSV files. For the presynaptic network map, we excluded inputs with less than 100 synapses. We created a weighted adjacency matrix in which each entry represented the total number of synapses from a given presynaptic neuron onto an antennal motor neuron. We used this matrix to construct a directed graph.

To quantify similarity in presynaptic input patterns across AMN pairs, we generated a weighted input matrix. Matrix entries reflected the total number of synapses from each presynaptic neuron onto that motor neuron pair, summed across all instances. Pairwise cosine similarity was computed between the input vectors of each motor neuron pair as the normalized dot product of their synapse-weighted input profiles, yielding a similarity score between 0 and 1 to reflect the degree of shared, synapse-weighted presynaptic input. We visualized this with a similarity matrix heatmap.

To verify the number of motor neurons entering the antennal nerves, we used the FAFB whole brain EM reconstruction to identify all annotated neurons in the antennal nerve. We then sorted these neurons by either area (nm^2^) or size (nm^3^) from largest to smallest to identify large cells innervating the antennae. While plotting the distribution of these neurons, we visually determined an area or size threshold to isolate large neurons, consistently revealing the same 7 pairs of large, efferent neurons in the two antennal nerves.

## Acknowledgements

This work was funded in part by grants awarded to M.P.S. from the National Institutes of Health Brain Initiative (R00NS114179 and U01NS131438) and the National Institute of Neurological Disorders and Stroke (R01NS140174). T.C.M. contributions to this material is based on work supported by the Air Force Office of Scientific Research under award number FA9550-25-C-B010 in the amount of $148,673. Igor Siwanowicz was supported through the Antibody Project Team at HHMI’s Janelia Research Campus for this work. Stocks obtained from the Bloomington Drosophila Stock Center (NIH P40OD018537) and the Korea Drosophila Resource Center (KDRC) were used in this study. Confocal experiments were performed in part with the Vanderbilt Cell Imaging Shared Resource (supported by NIH grants S10OD021630, CA68485, DK20593, DK58404, DK59637, and EY08126). The authors would like to thank Lisa Fenk for helpful discussion and suggesting driver lines labeling both eye and antennal motor neurons, and Bradley Dickerson for helpful comments on the manuscript.

---

**Supplemental Figure 1.**
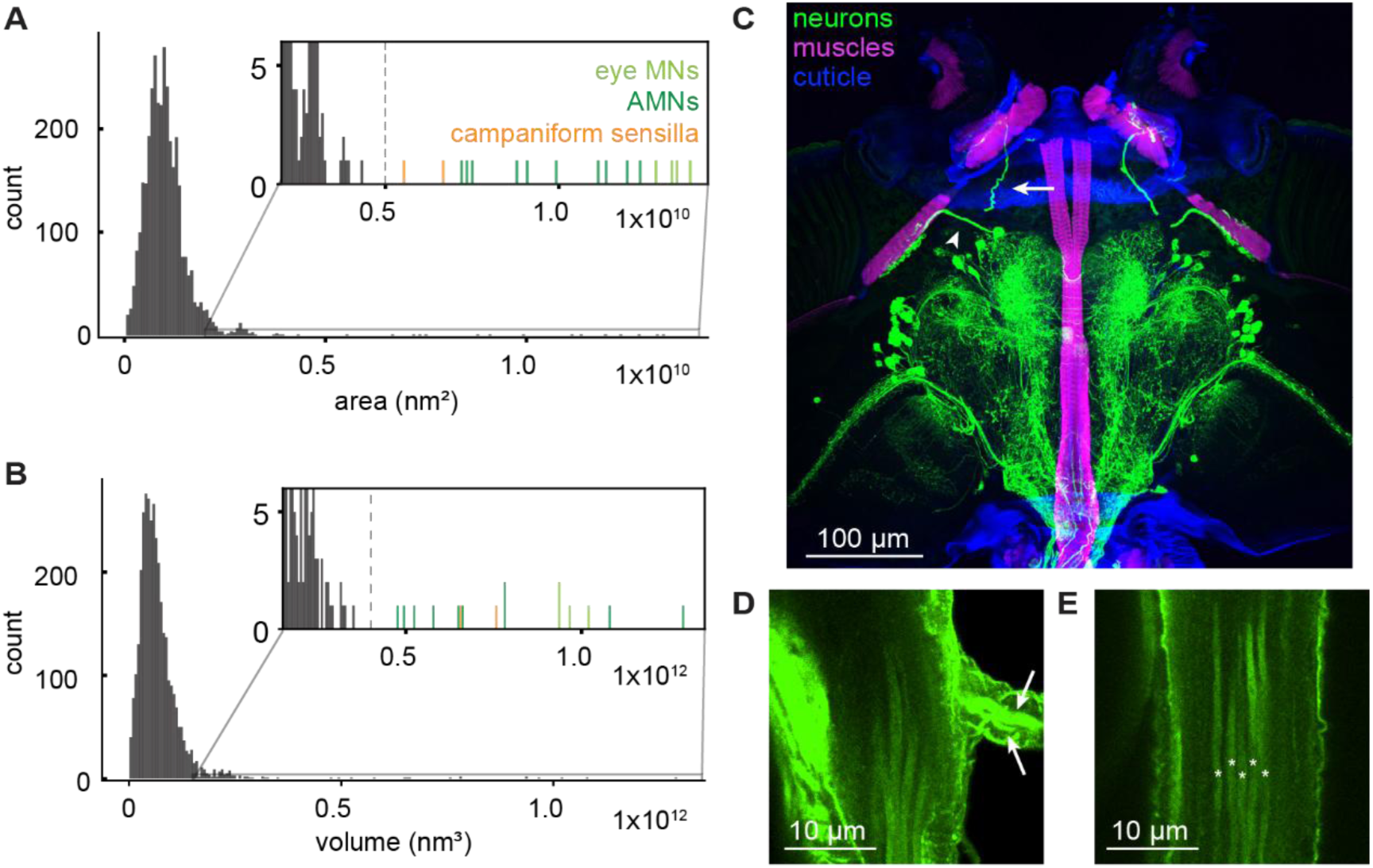
Seven motor neurons travel through the antennal nerve. (A) Histogram of axon cross-sectional areas (nm^2^) for all neurons in the antennal nerve from the FAFB data set. Inset: eye motor neurons (eye MNs, light green), antennal motor neurons (AMNs, dark green), and the campaniform sensilla neurons (orange) are the largest cells in the antennal nerve by area. (B) Histogram of axonal volume (nm^3^) for all neurons in the antennal nerve. Inset: eye motor neurons, antennal motor neurons, and the campaniform sensilla neurons are the largest cells in the antennal nerve by volume. (C) Horizontal section of the *Drosophila* head showing eye and antennal motor neurons (green) traveling to their respective muscle targets (magenta). Cuticle is shown in blue. Eye motor neurons branch from the antennal nerve to innervate the retinal muscles (arrowhead points towards the left eye motor neuron axon), whereas antennal motor neurons continue through the nerve to innervate the antennal muscles (arrow points towards left antennal motor neuron axon). (D) Eye motor neurons leaving the antennal nerve through a proximal nerve branch (axons indicated by arrows). (E) Five large axons, each indicated by an asterisk, continue through the antennal nerve and ultimately into the antennal first segment.

**Supplemental Figure 2.**
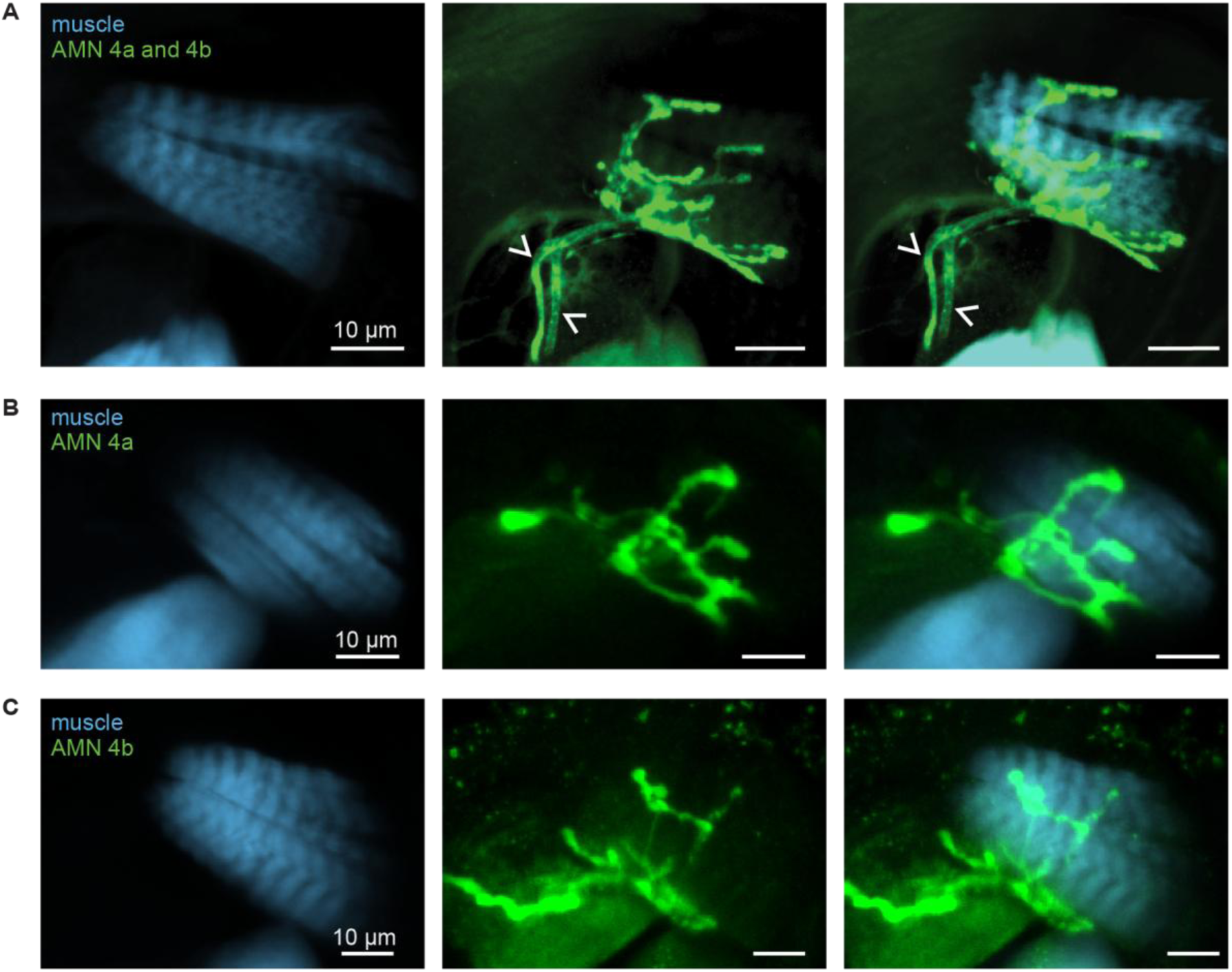
Anatomy of muscle 4 motor neurons. (A) Two motor neurons labeled by *R18D07 >10XUAS-IVS-myr::tdTomato* innervate muscle 4. Arrows indicate the two axonal projections. (B) AMN 4a innervates both muscle partitions, as labeled by *R18D07 > 20XUAS-SPARC2I Chrimson::tdTomato.* (C) AMN 4b innervates both muscle partitions, as labeled by *R75G05 > UAS-Chrimson*.

**Supplemental Figure 3:**
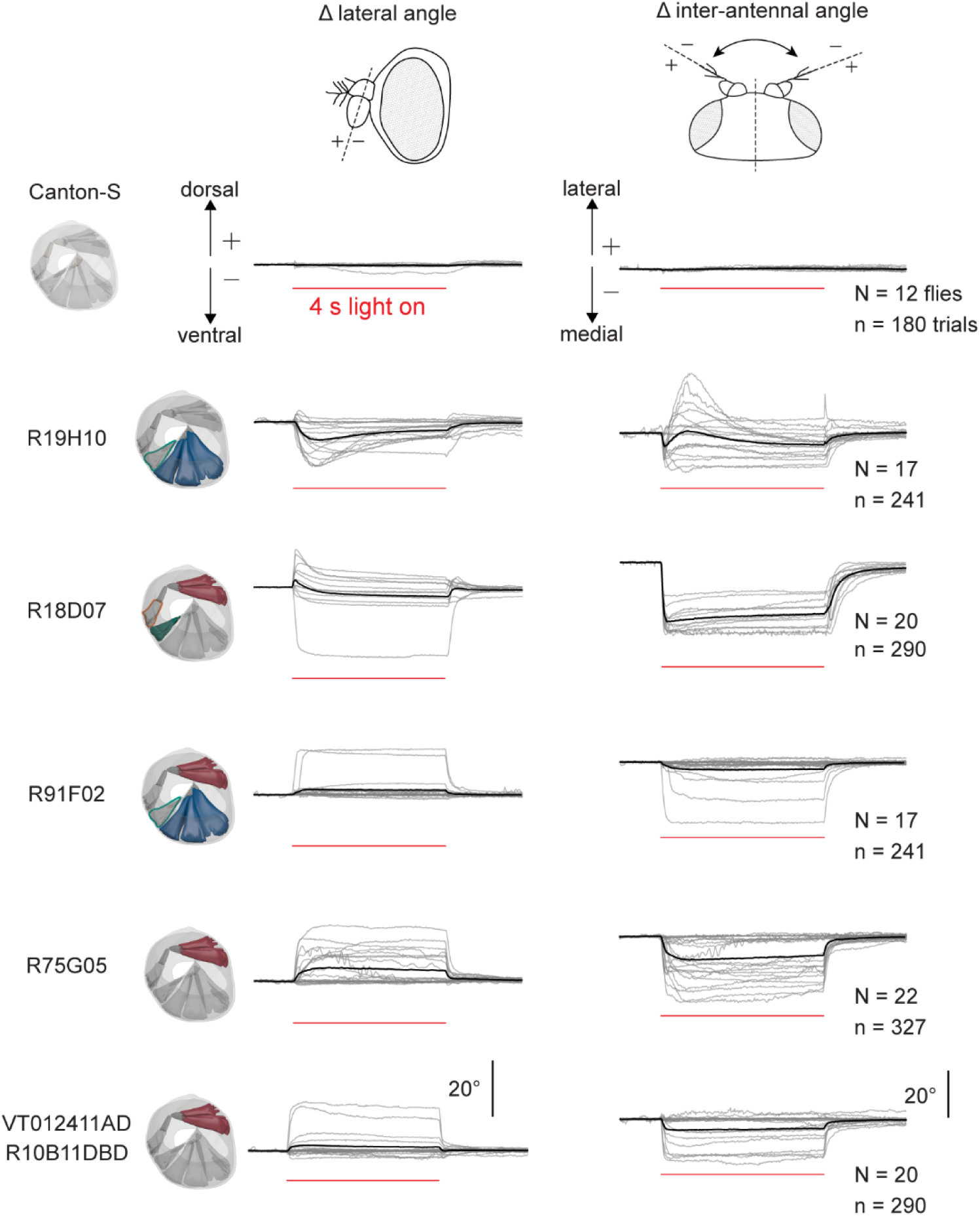
Activation of driver lines targeting subsets of antennal motor neurons. Left column: Schematics representing the targeted muscles for each driver line. Outlined muscles indicate stochastic effector expression in the associated motor neuron for some muscles. Center column: Antennal movements in response to the activation stimulus as tracked from the lateral camera. Downward antennal movements are negative, and upward movements are positive. Right column: Antennal movements in response to the activation stimulus as tracked from the dorsal camera. Movements toward the midline of the fly are negative and movements away from the midline are positive. Grey lines represent the average antennal response of individual flies and black traces represent the average response across flies.

## Supplemental methods

### Preparation for Supplemental Figures 1 and 2

Flies were anesthetized on ice and briefly washed with 70% ethanol. Heads were isolated and proboscises were removed under 2% paraformaldehyde/PBS/0,1% triton X-100 (PBT) and fixed in this solution for 4 h at 4°C. After washing in PBT, the samples were embedded in 7% agarose and sectioned on Leica Vibratome (VT1000s) horizontally at 0.25 mm. The slices were bleached in 10mg/ml sodium borohydride (NaBH4) in PBS, washed and blocked in PBS with 1% triton X-100 and 3% NGS (normal goat serum) for an hour at RT, followed by 48 h incubation in the same solution containing Texas Red-X Phalloidin (1:50, Life Technologies #T7471), rabbit anti-DsRed polyclonal antibodies (1:1000, TaKaRa Bio USA) and a chitin-binding dye Calcofluor White (0.1 mg/ml, Sigma-Aldrich #F3543-1G) at room temperature with agitation. After a series of three ∼30 minutes-long washes in PBT, the sections were incubated for another 24h in the above buffer containing secondary antibodies (1:1000, goat anti-rabbit, Thermo Fisher #A32731). The samples were then washed in PBT and fixed for 4 hours, RT in 2% paraformaldehyde to reduce leaching of bound phalloidin from muscles during the subsequent ethanol dehydration step. To avoid artifacts caused by osmotic shrinkage of soft tissue, samples were gradually dehydrated in glycerol (2-80%) and then ethanol (20 to 100 %; Ott 2008) and mounted in methyl salicylate between 2 #1 cover slips with custom-made spacers (Scotch office tape [2 layers = ∼120 µm] + double-sided tape [∼100 µm]) for imaging.

### Imaging and rendering

Serial optical sections were obtained at 1 µm intervals with a LD-LCI 25x/0.8 NA objective, or 0.3 µm with a 40x, 1.3 NA PlanApo objective on a Zeiss LSM 880 confocal microscope.

Samples were imaged using 405, 488 and 594 nm lasers, respectively. Blue-green emission of Calcofluor bleeds into the green channel; sequential mode of scanning was therefore applied.

## Notes

### Competing Interest Statement

The authors have declared no competing interest.

